# Polygenic prediction of school performance in children with and without psychiatric disorders

**DOI:** 10.1101/2020.07.15.203661

**Authors:** Veera M. Rajagopal, Betina B Trabjerg, Jakob Grove, Henriette T. Horsdal, Liselotte Petersen, Cynthia M. Bulik, Jonas Bybjerg-Grauholm, Marie Bækvad-Hansen, David M Hougaard, Ole Mors, Merete Nordentoft, Thomas Werge, Preben Bo Mortensen, Esben Agerbo, Anders D. Borglum, Ditte Demontis

## Abstract

Suboptimal school performance is often seen in children with psychiatric disorders and is influenced by both genetics and the environment. Educational attainment polygenic score (EA-PGS) has been shown to significantly predict school performance in the general population. Here we analyze the association of EA-PGS with school performance in 18,495 children with and 12,487, without one or more of six psychiatric disorders and show that variance explained in the school performance by the EA-PGS is substantially lower in children with attention deficit hyperactivity disorder (ADHD) and autism spectrum disorder (ASD). Accounting for parents’ socioeconomic status obliterated the variance difference between ADHD–but not ASD–and controls. Given that a large proportion of the prediction performance of EA-PGS originate from family environment, our findings hint that family environmental influences on school performance might differ between ADHD and controls; studying the same further will open new avenues to improve the school performance of children with ADHD.

## Introduction

The highest level of education attained by an individual, referred to as educational attainment (EA), strongly influences of a range of life outcomes including physical health, mental health, socioeconomic and behavioral outcomes^1^. Individuals differ widely in their EA. Much of these differences are due to genetic and environmental factors, and the interplay between the two^2^.

EA is moderately heritable^2^ and the heritability estimates vary widely; twin studies based estimate is around 40%^3^. Some of the largest GWASs (> 1 million) conducted to date include that of EA^4^ and they have demonstrated that EA is highly polygenic and thousands of variants together contribute to the heritability of EA. The effects of these variants can be combined in to a single score, referred to as polygenic score (PGS). EA-PGS has been shown to associate strongly with EA in independent samples, explaining up to 12.7% of the phenotypic variance in EA^4^ and up to 14% of the phenotypic variance in school grades^5-7^.

Although EA-PGS strongly associates with school grades, its association is influenced by numerous factors such as ancestry^8^, environmental factors^9^ and indirect genetic effects such as genetic nurture^10^ and social genetic effects^11^. Existing GWASs of EA have been conducted in individuals from European ancestries^4; 12; 13^. Hence, EA-PGS associates with EA poorly in individuals from non-European ancestries such as African-American ancestries^4; 14^. Several environmental factors such as socioeconomic status, access to education as well influence EA strongly and therefore the strength of the association of EA-PGS with EA reduces when variations in such environmental factors increase^15^. Moreover, the strength of the association of EA-PGS with EA drops substantially when analyzing siblings than when analyzing unrelated individuals^16^, indicating that a large proportion of EA-PGS’s association with EA is mediated through family environment.

Individuals with—or those at risk for—psychiatric disorders often score lower than individuals without psychiatric disorders in school^17^. Although genetic factors related to cognition partly explain this association^18^, other factors such as family socioeconomic status (SES) contribute as well, for example, individuals with schizophrenia and ADHD are often brought up in families with low SES compared to individuals from the general population^19-21^. Additionally, there is a significant sharing of genetic risk variants between psychiatric disorders and cognitive phenotypes such as EA^22^ and intelligence^23^. Given these backgrounds, it can be hypothesized that the EA-PGS might influence EA differently in individuals with psychiatric disorders than in the general population. However, this has not been evaluated so far.

The aim of the current study is to evaluate if the association between EA-PGS and school performance differ between individuals with and without psychiatric disorders. To do this, we studied the association of EA-PGS with school performance in 30,982 individuals from the Danish iPSYCH and ANGI cohorts^24; 25^, 60% of whom were diagnosed with one or more of six major psychiatric disorders—ADHD, ASD, schizophrenia (SCZ), bipolar disorder (BD), major depressive disorder (MDD) and anorexia nervosa (AN). Through an analysis involving the full cohort we first confirmed that EA-PGS strongly associate with school performance. Then we proceeded with subgroup analysis where we tested for differences in the association of EA-PGS with school performance across six psychiatric groups and controls (i.e. those without any of the six disorders). Additionally, we also explored the differences in the association of some of the known environmental predictors of school grades (parental education and employment status) across the six psychiatric disorder groups and controls. We then compared the results with that of EA-PGS to infer how genetic and environmental influences of school performance differed across the six psychiatric disorder groups and controls.

## Results

### Sample characteristics

Our study individuals comprised of 18,495 with one or more of six major psychiatric disorders (ADHD=5238, ASD=3859, SCZ=1356, MDD=10,123, BPD=839 and AN=1680) and 12,487 without any of these six disorders^22^ (Table 1). We assessed the school performance based on six different school grades (Danish written, oral and grammar, English oral, and mathematics written and oral, or problem solving with and without aids; Methods). The school grades were from the exit exam given at the end of compulsory schooling in Denmark. The students were on average 15.7 years old (SD=0.42) when they sat for their exit exams. It is important to note that among those with psychiatric disorders we studied, many received their psychiatric diagnoses only after they sat for the exams (Table 1). Yet their school performance differed significantly from the controls, which we have reported previously^22^. Hence, we considered everyone with a psychiatric diagnosis in the register (as of Dec 2016) as cases irrespective of when they received their diagnoses. The sample characteristics within each disorder group and controls are summarized in Table 1.

**Table 1.** Sample characteristics.

| Variable | Schizophrenia | Bipolar disorder | MDD | ADHD | ASD | Anorexia | Controls |
| --- | --- | --- | --- | --- | --- | --- | --- |
| Sample size | 1356 | 839 | 9719 | 5238 | 3859 | 1680 | 12487 |
| Females | 668 (49.3%) | 561 (66.9%) | 7066 (69.8%) | 1748 (33.4%) | 1017 (26.4%) | 1568 (93.3%) | 6327 (50.7%) |
| Age on Dec 2016 | 26.1 (2.9) | 26.8 (2.8) | 26.2 (3.3) | 22.9 (4.1) | 22.1 (3.9) | 24.8 (3.7) | 23.4 (4.2) |
| Exam age<sup>1</sup> | 15.7 (0.4) | 15.6 (0.4) | 15.6 (0.4) | 15.8 (0.4) | 15.8 (0.5) | 15.6 (0.4) | 15.6 (0.4) |
| Age at first diagnosis<sup>2</sup> | 21.15 (2.9) | 21.86 (3.2) | 19.0 (3.3) | 14.8 (5.6) | 12.8 (5.0) | 16.3 (3.3) | - |
| Diagnosed before exam<sup>3</sup> | 38 (2.8%) | 17 (2.1%) | 1668 (16.5%) | 3025 (57.8%) | 2880 (74.6%) | 819 (48.9%) | - |
| Mean school grades<sup>4</sup> | 6.0 (2.2) | 6.7 (2.3) | 6.4 (2.3) | 5.3 (2.2) | 6.7 (2.3) | 8.0 (2.3) | 6.9 (2.4) |

The value represents either frequency (proportion) if categorical variable or mean (SD) if continuous variable.
<sup>1</sup>Exam age is the age of the individuals when they sat for the exit exam.
<sup>2</sup>The age at first diagnosis was calculated based on the first time the diagnosis was recorded in the register.
<sup>3</sup>The date of first diagnosis and the date of exit exam were compared to identify if the individual has received the diagnosis before they sat for the exam.
<sup>4</sup>The mean value was calculated across all the grades: Danish written, oral and grammar and English oral and mathematics written and oral (if sat for the exam before on or before 2006) or problem solving (if sat for the exam after 2006); The lowest and highest mean values observed were -1.7 and 12.

### Derivation of a proxy variable for overall school performance

All the six grades that we analyzed were strongly correlated with each other suggesting that those who performed well in one subject performed well in others as well^22^ (Figure 1a). To measure the overall performance, we therefore did a principal component analysis (PCA) and extracted the first principal component, PC1 (hereafter, E1), that explained maximum variance in all the subject grades (Figure 1b). This is equivalent to extracting a general cognitive ability factor (g) from a PCA of multiple cognitive test scores^26^. We have previously reported the genetic architecture of E1 by performing a genome-wide association study (GWAS)^26^ and showed that there is an almost complete genetic overlap between E1 and EA (r_g_=0.90; SE=0.01; P=4.8×10^−198^). Hence, here we used E1 as a proxy phenotype for EA and studied its association with EA-PGS across the six psychiatric disorder groups and controls.

**Figure 1.**
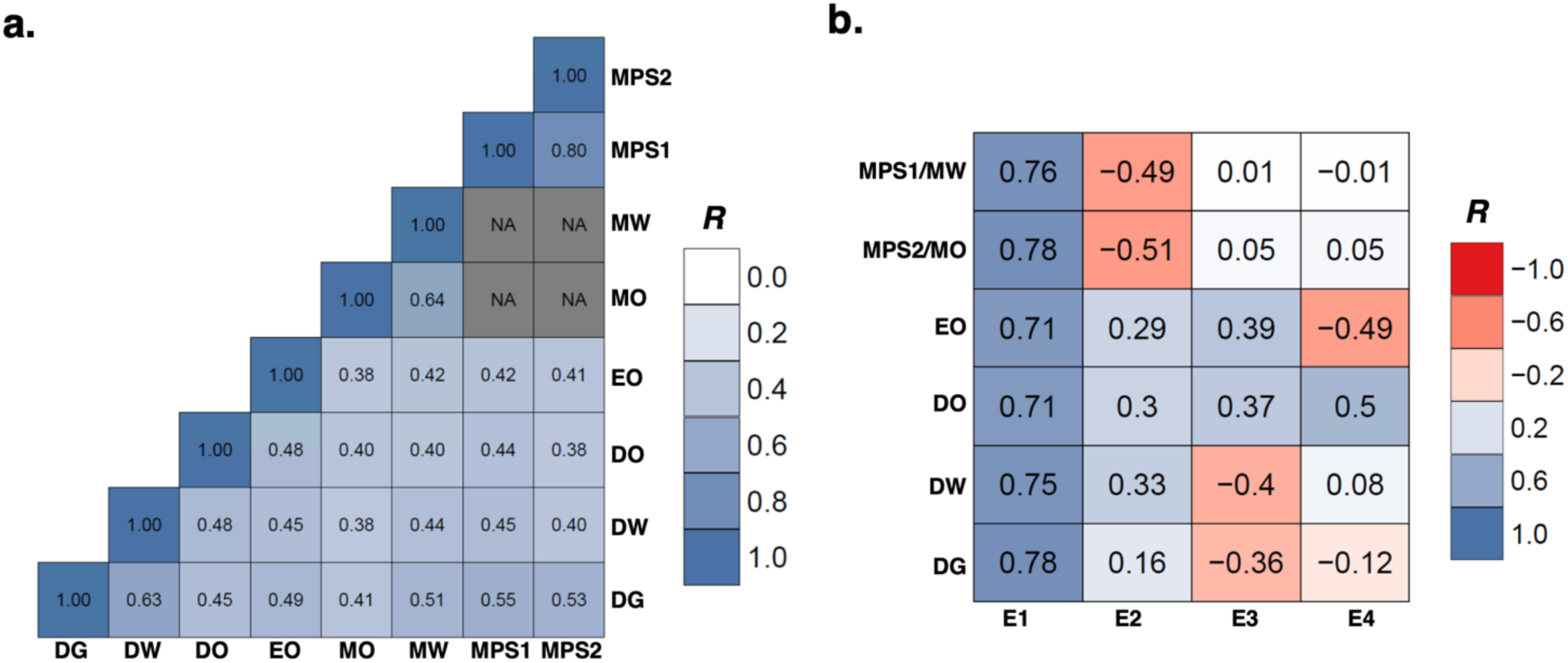
Correlations within subject-grades and between subject-grades and principal components. **a**. Heatmap of pairwise correlations across the subject-grades in the full sample (N=30,982); the values correspond to Pearson’s correlation coefficient; NA values mean that there is no overlapping samples for the subject pairs; **b**. Heatmap of correlations between the first four principal components and subject-grades in the full sample (N=30,982); the value correspond to Pearson’s correlation coefficient; DG – Danish grammar; DW – Danish written; DO – Danish oral; EO – English oral; MO – Math oral; MW – Math written; MPS1 – Math problem solving 1 (without aids); MPS2 – Math problem solving 2 (with aids, e.g. calculator)

### Overall association of EA-PGS with E1

First, we studied the association of EA-PGS with E1 in the full sample (N=30,982) that included both individuals with and without psychiatric disorders. We constructed EA-PGS for our study individuals at ten P value thresholds using the variant effect sizes reported in the recent largest GWAS of EA^4^ (Methods). We included exam age (age at the time of examination), sex and psychiatric diagnoses as covariates along with others in the analyses (Methods). EA-PGS at all ten thresholds associated significantly with E1 and explained substantial proportion of variances in E1 (Supplementary Table 1). EA-PGS at threshold P<0.05 explained the maximum variance (R^2^=8.4%) and so we used this threshold for further analysis.

**Figure 2.**
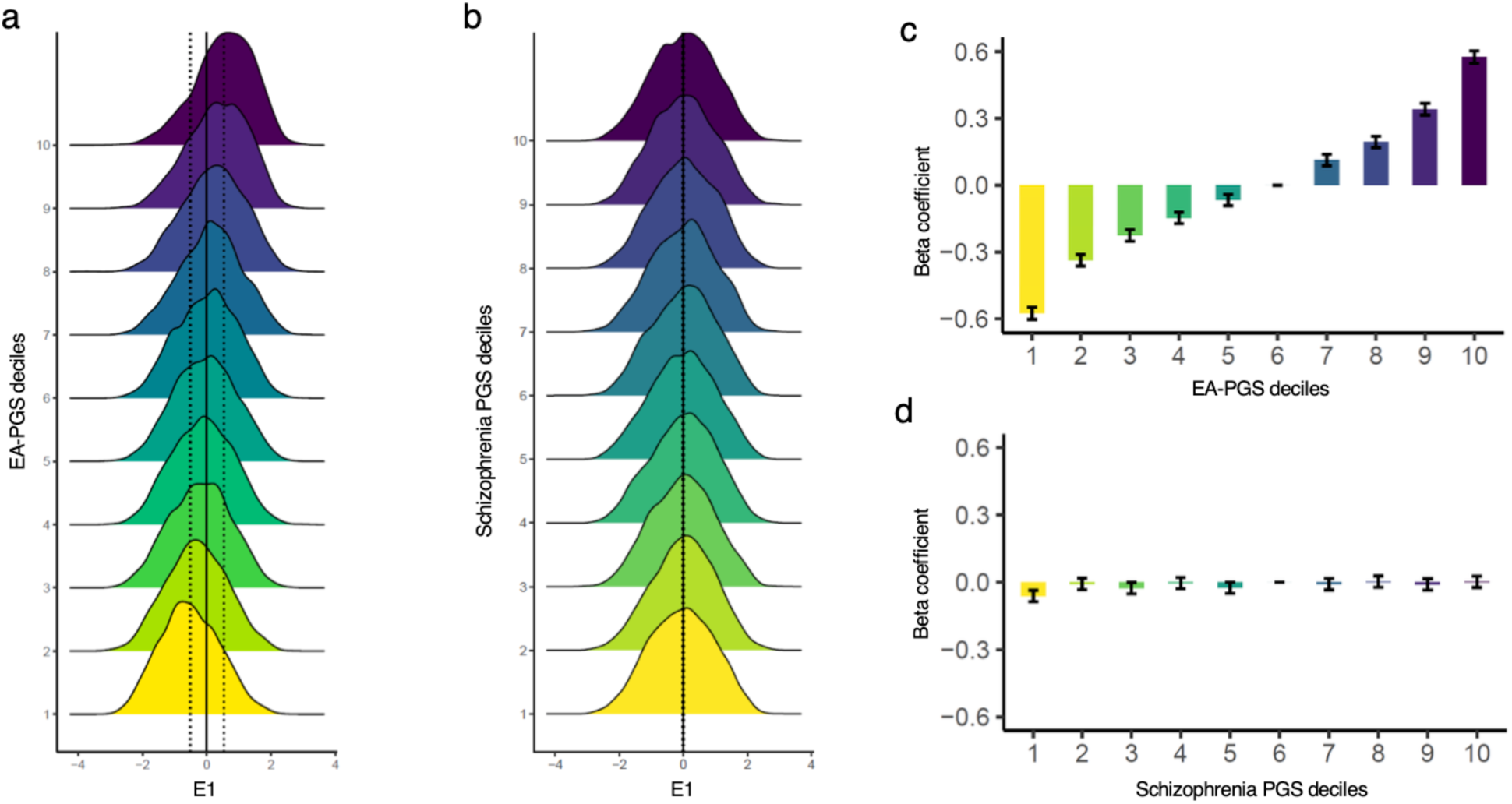
Distribution of E1 across PGS deciles. **a**, Distribution of E1 across deciles based on EA-PGS. The E1 scores are residualized for all the covariates and then the residuals are plotted. Centre solid line correspond to full sample mean. The left and right dotted lines correspond to means of the first and last deciles respectively. **b**, Same as a, but the deciles are created using schizophrenia PGS as a negative control. **c**, E1 scores of each decile are compared against decile 6 using logistic regression adjusted for all covariates; the betas and the standard errors are plotted. **d**, same as c, but the deciles are created using schizophrenia PGS as a negative control.

To further explore the association of EA-PGS with E1, we performed a decile analysis. We divided the entire sample into deciles based on EA-PGS and then explored how the E1 mean scores trended from the lowest decile to the highest (Figure 2). The E1 scores were residualized for all the covariates and then Z-normalized to have mean zero and standard deviation (SD) one in the whole sample. The E1 mean was lowest for decile one (mean=-0.51; SD=0.92), which gradually increased through decile two until decile ten, which had the highest E1 mean (0.53; SD=0.92; Figure 2a; Supplementary Table 2a). A linear trend was observed reflecting a strong correlation between EA-PGS and E1. The E1 mean values of deciles one to five were significantly lower than zero (i.e. the mean of the full sample, which can be assumed as the population mean) and the E1 mean values of deciles seven to ten were significantly higher than zero (Figure 2a; Supplementary Table 2a). The E1 mean value of decile six was not significantly different from zero and so decile six can be considered representative of the population. To quantify the effect sizes of the individual deciles, we compared the E1 values of each of the deciles one to five and deciles seven to ten against decile six using logistic regression adjusted for all the covariates. The beta values from the regression analyses trended uniformly across the deciles similar to how the E1 mean values did (Figure 2c; Supplementary Table 2b). As a negative control, we repeated the above analyses using schizophrenia PGS as schizophrenia had a minimal genetic correlation with E1 (r_g_=0.06; SE=0.03; P=0.06;). This was performed to show how the distribution of effect sizes look if the deciles were created based on a PGS that associated weakly with E1 (Figure 2b, 2d; Supplementary Table 2a, 2b).

We repeated the decile analysis using only the controls and found similar results except that the effect sizes were slightly larger (Fig. 2—figure supplement 1; Supplementary Table 3). Hence, including individuals with psychiatric disorders has slightly attenuated the effect sizes suggesting that the association of EA-PGS with E1 may not be as much stronger in individuals with psychiatric disorders as they are in the general population. In summary, our initial analysis confirmed that EA-PGS significantly associate with school performance and explain a substantial proportion of variance in school grades thereby corroborating with previous studies^5-7^.

### Benchmarking effect sizes of polygenic deciles against ADHD

Given the strong correlation between EA-PGS and school performance, it can be expected that school performance of those at the lower end of the EA-PGS distribution might be comparable to the school performance of those with learning difficulties. However, to our knowledge, this has not been evaluated so far using a proper reference group such as those with learning difficulties. In our cohort, individuals diagnosed with ADHD scored the lowest (on average) of all the psychiatric disorder groups and controls suggesting that many of these individuals might have learning difficulties^22^. It has been also well documented that individuals with ADHD often experience learning difficulties in school^27-29^. Hence, we benchmarked the effect sizes of EA-PGS deciles by using the individuals with ADHD as the reference group.

We grouped the controls into deciles based on their EA-PGS. Then we combined the controls and those with ADHD and Z-normalized their E1 (after residualizing for covariates) to have mean zero and SD one in the combined sample. Then we compared the mean value of E1 in each of the ten control deciles with that of ADHD (Figure 3; Supplementary Table 4). Except the lowest decile, the mean E1 of the rest of the deciles (two to ten) were significantly higher than the mean E1 of ADHD (Figure 3a; Supplementary Table 4a). The mean E1 of decile one (−0.37; SD=0.94) was similar to the mean E1 of ADHD (−0.41; SD=0.92; P=0.19) suggesting that the school performance of those in the lowest polygenic decile was similar to that of ADHD. We also compared the E1 of each of the deciles against ADHD using logistic regression analysis adjusting for all the covariates and found similar results (Figure 3b; Supplementary Table 4b). In summary, our analysis suggested that even among those without any apparent learning difficulties, EA-PGS was able to identify a group of individuals whose school performance was comparable to the school performance of those with ADHD, who were known to often experience learning difficulties.

**Figure 3.**
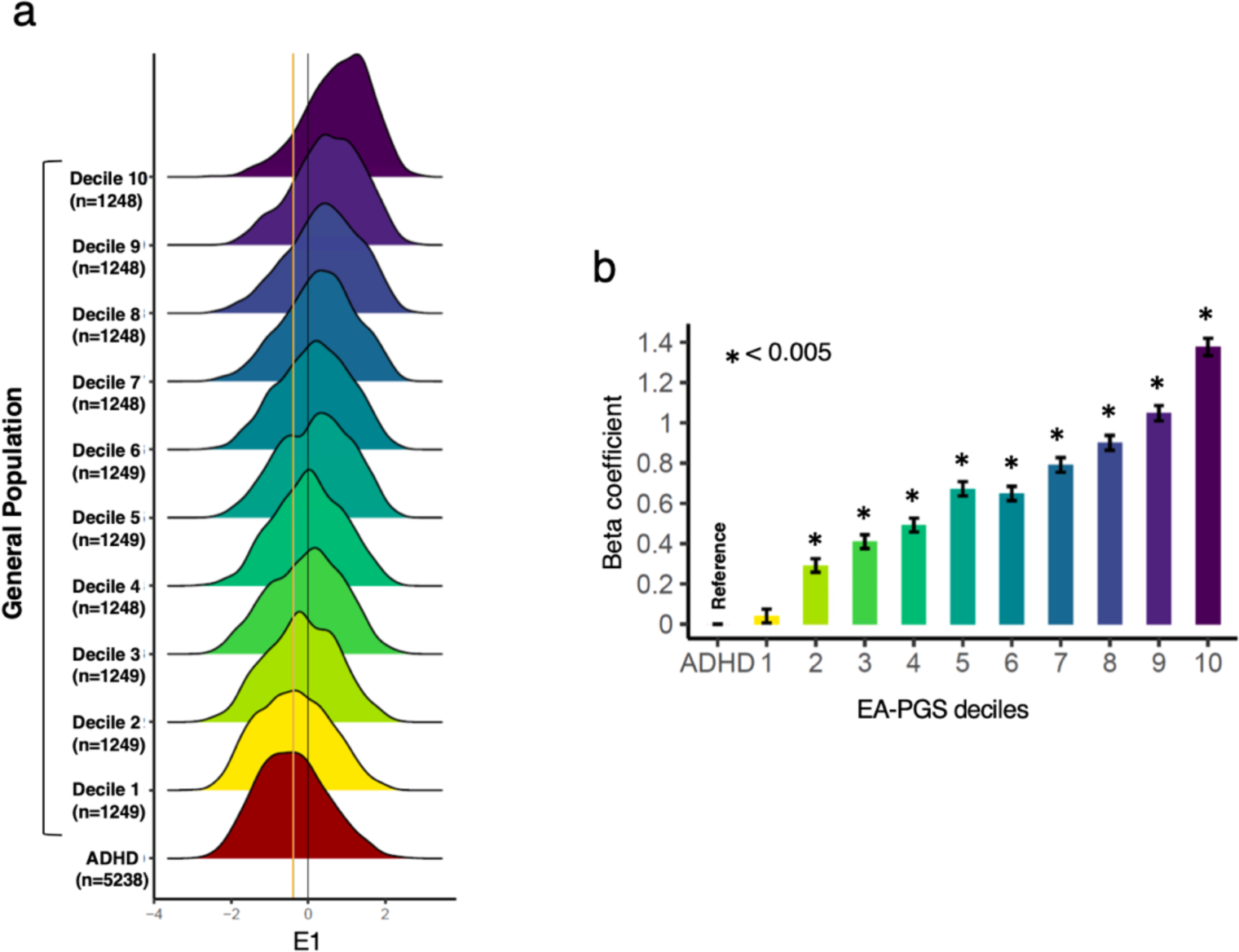
Benchmarking polygenic deciles against ADHD. **a**, Distribution of E1 across deciles based on EA-PGS (only controls; N=12,487) and ADHD (N=5,238). The E1 scores are residualized for all the covariates and then the residuals are plotted. The center black line corresponds to full sample mean. The means of decile 1 and ADHD are shown orange and blue lines respectively. **b**, E1 scores in each decile is compared against ADHD using logistic regression adjusted for covariates (methods). Beta and standard errors are plotted.

### Psychiatric disorder specific association of EA-PGS with E1

Given that we have confirmed that EA-PGS strongly associate with E1, we next studied how the association of EA-PGS with E1 varied across the psychiatric disorder groups and controls. We performed association analysis within each psychiatric group and controls using linear regression adjusted for all the covariates (same as the ones used in the main analysis, except psychiatric diagnoses). The strength of the associations was assessed based on the proportion of variance in E1 explained by EA-PGS (incremental R^2^; Methods). We estimated standard errors for R^2^ by bootstrapping 1000 times and therefore we were able to statistically test if R^2^ in each of the psychiatric disorder group differed significantly from R^2^ in the controls (Methods).

The distributions of E1 and EA-PGS across the disorder groups and controls are shown in Figure 4a and summarized in Supplementary Table 5. Briefly, the E1 mean values of ADHD, SCZ and MDD were significantly lower than that of controls and the E1 mean values of ASD, BD and AN were significantly higher than that of controls. The E1 distributions were largely similar across the groups. While the phenotypic variances of E1 in MDD, BD and AN did not differ significantly from that in the controls, the phenotypic variances of E1 in ADHD, ASD and SCZ were significantly lower than in the controls (but the differences were only modest).

**Figure 4.**
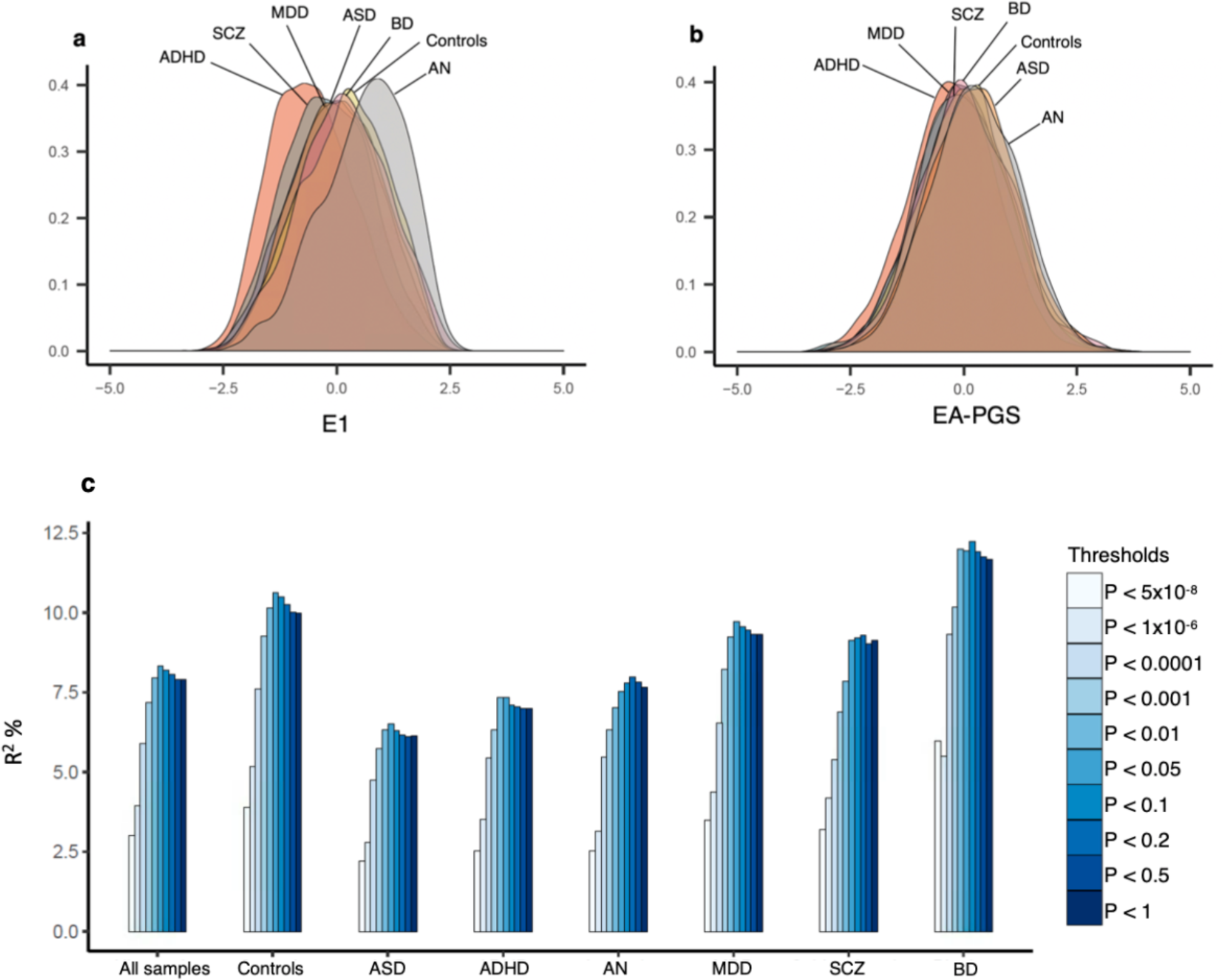
Polygenic prediction of school grades in controls and psychiatric disorders. **a**. Distribution of E1 across groups; **b**. Distribution of EA-PGS across groups; both E1 and EA-PGS were Z-normalized to have mean zero and SD one in the full sample (N=30,982); **c**. Variances explained by EA-PGS constructed at ten different P value thresholds. For sample size for each group, refer to Table 1. The variance explained is measured as incremental R^2^ (methods).

The distribution of EA-PGS across psychiatric disorders and controls are shown in Figure 4b and summarized in Supplementary Table 5. Briefly, the EA-PGS of all the psychiatric disorder groups except BD, differed significantly from controls. The mean EA-PGS of ADHD, MDD and SCZ were lower than controls and that of ASD and AN were higher than controls. The variance in the EA-PGS in any of the disorder groups did not differ significantly from controls.

Stratified polygenic score analyses revealed wide differences in the association of EA-PGS with E1 across the six psychiatric disorder groups and controls (Figure 4; Supplementary Table 6). We computed statistical difference in R^2^ between cases and controls at P value thresholds that corresponded to the maximum variance explained (Figure 5a). Firstly, EA-PGS explained more variance in E1 in controls than in the full sample; *R*^*2*^ in the controls was 10.61%, which was 1.28 times higher than *R*^*2*^ in the full sample (P_diff_=1.9×10^−5^). Secondly, EA-PGS explained substantially lower variances in E1 in ASD and ADHD than in the controls; R^2^ in ASD and ADHD were 6.52% and 7.3%, which were 38.7% and 31.1% less respectively compared to R^2^ in the controls and the differences were statistically significant (ASD: P_diff_=2.1×10^−6^; ADHD: P_diff_=2.9×10^−5^). Thirdly, EA-PGS explained higher variance in E1 in BPD (R^2^=12.2%; P_diff_=0.41) and lower variance in E1 in AN (R^2^=7.9%; P_diff_=0.15) than in the controls. However, the differences were not statistically significant, which might be due to that the sample sizes in these groups were not as large as the sample sizes in ASD and ADHD. Lastly, the variances explained in E1 in SCZ (R^2^=9.2%; P_diff_=0.21) and MDD (R^2^=9.4%; P_diff_=0.20) were almost similar to that in controls. Since many in the ADHD group were comorbid for ASD and vice versa, we repeated the analysis in individuals with only ADHD and only ASD and found similar results (Supplementary Table 6). Hence, the low variance in ADHD group was not due to a subset of individuals who were comorbid with ASD and likewise the low variance in ASD group was not due to a subset of individuals who were comorbid with ADHD. Compared to other psychiatric disorders, ASD is relatively more heterogenous constituting of subtypes that differ significantly in terms of cognitive functions^30^. Hence, we also tested if the variance explained is low across subtypes of ASD. We grouped the ASD individuals into three groups: Asperger’s (ICD-10 F84.5); miscellaneous pervasive developmental disorders (PDD), which included other PDD and PDD unspecified (F84.9 and F84.8); other subtypes, which included rest of the subtypes including childhood autism and atypical autism (as there were only very few in each of the other ASD subtypes, we clumped them together into a single group). We repeated the analysis in the three groups separately (Supplementary Table 6). Among the three groups, the R^2^ was lowest in Asperger’s (R^2^=5.4%; P_diff_=4.4×10^−7^) and was significantly different from the R^2^ in controls. The R^2^ in other two groups were lower, but did not differ significantly from the R^2^ in controls (miscellaneous PDD: R^2^=8%, P_diff_=0.04, adjusted P_diff_=0.13; other subtypes: R^2^=6.7%; P_diff_=0.03; adjusted P_diff_=0.11). This could be partly due to that the sample size of these two groups were lower compared to sample size in Asperger’s.

**Figure 5.**
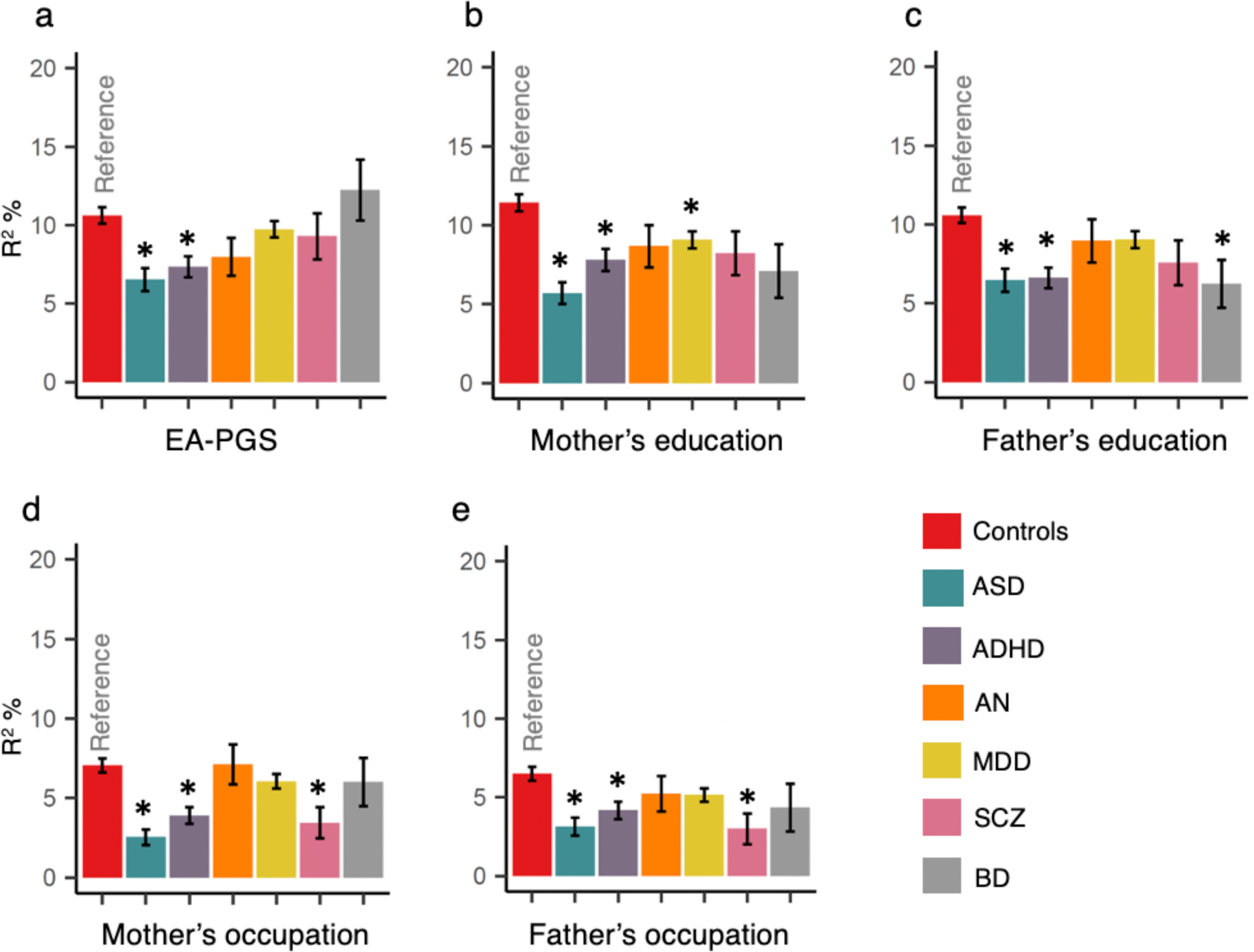
Variance explained in E1 in controls and in psychiatric disorders by. **a**, EA-PGS constructed using threshold that explained the maximum variance in the respective groups; **b**, Mother’s education status; **c**, Father’s education status; **d**, Mother’s employment level; e, Father’s employment level. Association analyses were performed using linear regression adjusted for covariates (see methods). The variance explained is assessed using incremental R2 (see Methods), which are shown as bars. The error bars represent standard errors estimates using bootstrapping. Star symbol indicates that the R2 is statistically different from the R2 in controls, assessed using Z test.

In summary, we found that the association between EA-PGS and E1 was not as much stronger in individuals with psychiatric disorder as it was in controls; although the associations in all the six disorders differed at least to some extent from that of the controls, a large and statistically significant difference was observed only in ASD and ADHD.

It is important to note that the low variance explained by the EA-PGS in E1 in ASD and ADHD compared to controls is not due to that the phenotypic variance in E1 is itself lower in ASD and ADHD compared to controls. We show this by creating two down-sampled subsets of controls (methods), one with matching phenotypic E1 variance with ASD and the other, with ADHD and demonstrating that the variance explained by the EA-PGS in E1 is still significantly lower in ASD and ADHD compared to the matched controls. To down-sample the controls, we iteratively removed one individual at a time from the controls and tested if after removing the individual the E1 variance reduced. If so, we excluded that individual, otherwise we put the individual back in the controls. We repeated this in a loop until the E1 variance of the controls matched with ASD or ADHD. For ADHD, we removed 431 individuals from the controls and the E1 variance in the controls dropped to 0.829, which matched with E1 variance in ADHD (0.827, variance difference P=0.90). When we repeated the PGS analysis, we found that the EA-PGS explained 9.72% of variance in E1 in controls, which was still significantly higher than ADHD (R^2^=7.3%; P_diff_=0.002). For ASD, we removed 168 individuals from the controls and the E1 variance in the controls dropped to 0.90, which matched with the E1 variance in ASD (0.90; variance difference P=0.95). When we repeated the PGS analysis, we found that the EA-PGS explained 10.2% variance in E1 in controls, which was still significantly higher than ASD (R^2^=6.5%; P_diff_=1.8×10^−5^).

### Psychiatric disorder specific association of socioeconomic factors with E1

Studies have shown that socioeconomic status (SES) of individuals with psychiatric disorders differ significantly from general population^19; 31^. Also, SES strongly influences school performance^32^. Hence, we hypothesized that EA-PGS had a weaker influence on school performance in individuals with psychiatric disorders compared to controls because environmental factors such as SES might have had a stronger influence. Also, it is known that as environmental influences on EA increase, genetic influences on EA decrease^15; 33; 34^. To test our hypothesis, we studied four SES indicators namely mother’s education and employment levels and father’s education and employment levels. All the four SES indicators differed substantially between psychiatric disorders and controls. Overall, the SES of ADHD, MDD, SCZ and BD were significantly lower and the SES of AN was significantly higher compared to the SES of controls (Supplementary Table 7). We studied the association of E1 with four SES indicators using linear regression adjusted for exam age, sex and PCA group (methods). The strength of the association was assessed based on the proportion of variance in E1 explained by the SES indicators (incremental R^2^; Methods). Similar to the PGS analysis, we estimated standard errors for R^2^ via bootstrapping in order to statistically compare R^2^ between groups. The results were opposite to what we expected: similar to the EA-PGS, SES too explained lower variance in E1 in individuals with psychiatric disorders compared to controls (Fig. 5; Supplementary Table 8). Specifically, the variances explained in E1 in ASD and ADHD were significantly lower than controls and the R^2^ differences were statistically significant with regard to all the four SES indicators. In other psychiatric disorders, only few of the comparisons were statistically significant: mother’s and father’s occupation explained significantly lower variance in E1 in SCZ than in the controls; mother’s education explained significantly lower variance in E1 in AN than in the controls; father’s education explained significantly lower variance in E1 in BD than in the controls (Fig 5; Supplementary Table 8).

In summary, the results suggested that the decrease in the influence of EA-PGS on school performance in individuals with ASD and ADHD is not due to a corresponding increase in the influence of SES on school performance in these individuals. On the contrary, the influence of SES on school performance was also lower in ASD and ADHD.

It is important to note that the SES difference between the groups itself is less likely to have affected the EA-PGS’s influence on school performance. For example, the SES of both ADHD and MDD groups were significantly lower than controls. However, the R^2^ was low only in ADHD, but not in MDD. In addition, when we divided the controls into five quantiles based on their SES and performed PGS analysis in each of the quantiles separately, we did not observe a significant difference in the R^2^ across the quantiles (Fig 5—figure supplement 1; Supplementary Table 9).

### Psychiatric disorder specific association of EA-PGS with E1 after adjusting for socio-economic factors

Given that both SES and EA-PGS both had weaker associations with school performance in individuals with psychiatric disorders than in the controls, it is likely that the attenuation of the association is caused by a factor that is common to both SES and EA-PGS. If so, adjusting for SES in the PGS analysis will attenuate the R^2^ differences between controls and psychiatric disorders (as the common factor gets cancelled out). So, we repeated the PGS analysis by including the four SES indicators as covariates. As we expected, after accounting for the SES, the R^2^ differences between controls and psychiatric disorders attenuated, but not in all the disorders (Fig. 6; Supplementary Table 10). The attenuation was largest for ADHD; before adjustment ADHD R^2^ was 31.1% lesser than controls R^2^, but after adjustment ADHD R^2^ was only 8.5% lesser than controls R^2^ and the R^2^ difference between ADHD and controls was no longer statistically significant (P_diff_=0.48). The attenuation was minimal in ASD; before adjustment ASD R^2^ was 38.7% lesser than controls R2 and after adjustment ASD R2 was 33% lesser than controls R2 and the R^2^ difference between ASD and controls remained statistically significant (P_diff_=0.005). In AN, almost no attenuation was seen; before adjustment AN R^2^ was 25.5% lesser than controls R^2^ and after adjustment AN R^2^ was 24.4% lesser than controls R^2^ and the R^2^ difference between AN and controls was not statistically significant either before or after adjustment (P_diff_=0.15). In MDD and SCZ, the R^2^ differences between the controls and these disorders almost fully disappeared after adjustment (MDD: P_diff_=0.98; SCZ: P_diff_=0.93). Unlike other disorders, we observed a marked increase in the R^2^ difference between BD and controls; before adjustment BD R^2^ was 15% higher than controls R^2^ and after adjustment BD R^2^ was 46.7% higher than controls R^2^; yet the R^2^ difference did not reach statistical significance (P_diff_=0.10; likely to be due to small sample size of BD).

**Figure 6.**
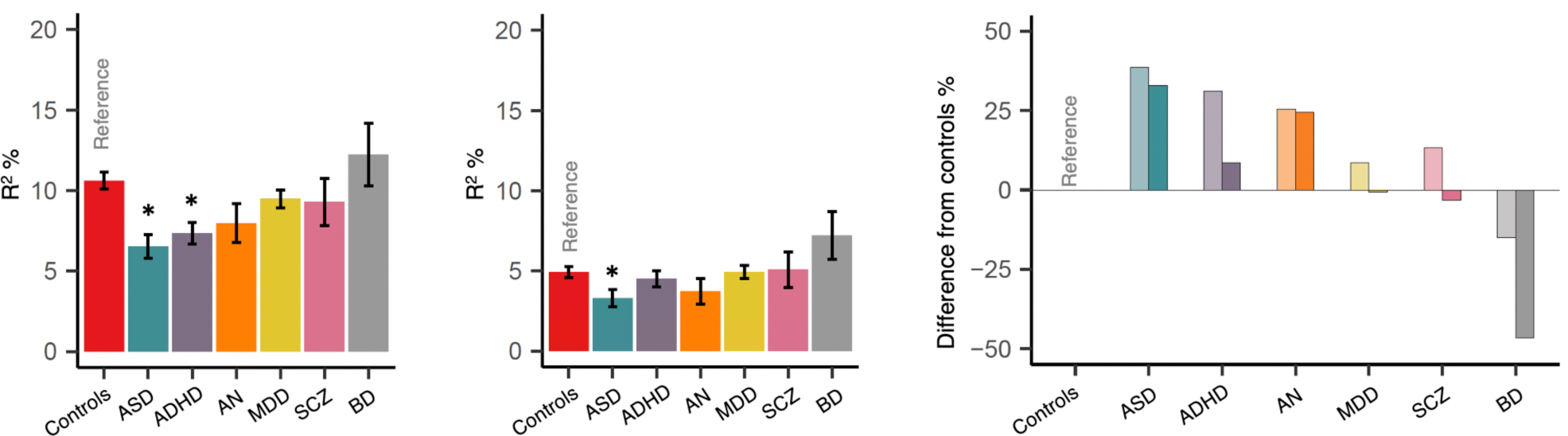
Association of EA-PGS after adjusting for SES. **a**, Association of EA-PGS with E1 before adjusting for SES (showed for comparison) **b**, Association of EA-PGS with E1 after adjusting for SES performed using linear regression adjusted for all the covariates used in the main analysis plus the four SES factors (father’s and mother’s education, father’s and mother’s employment); R2-Variance explained in E1 by EA-PGS; Error bars represent standard errors calculated using bootstrapping; Star symbol indicates the R2 is statistically different from R2 in controls **c**, Percentage difference in R2 between psychiatric groups and controls before (lighter shades) and after (darker shades) SES adjustment. Negative values indicate that the R2 in the psychiatric group is higher than the R2 in controls.

In summary, the large R^2^ difference between ADHD and controls reduced substantially after adjusting for SES, but the R^2^ difference between ASD and controls did not. Hence, the weaker association of EA-PGS with E1 in ADHD compared to controls is mainly due to a factor that is common between EA-PGS and SES. In ASD, however, this is not the case, hence the weaker association of EA-PGS with E1 in ASD compared to controls might be due to other sources of variation influencing E1.

## Discussion

Most of the previous studies on the association of EA-PGS with EA or school performance were based on individuals from the general population^5-7^. It was therefore not clear if results from such studies could be generalized to those with psychiatric disorders. Nation-wide register based cohorts such as iPSYCH^24^ are well suited to address such research questions as they offer a unique opportunity to study both the genetic and environmental factors of multiple psychiatric disorders at the same time within a single population setting. Also, currently iPSYCH has the largest sample size with information on both school grades and genotypes^22^. Hence, we studied the association of EA-PGS with school performance in iPSYCH and focused specifically on how the associations differed between those with and without psychiatric disorders.

As we had information on multiple types of school grades for each individual, we were able to perform a PCA and capture the overall school performance as a latent factor E1^22^. Defining the phenotype this way is better than taking an average across all the grades as latent variable captures variance across all the subjects more uniformly. For example, a high E1 score in an individual indicates that the individual performed equally well in all the subjects.

We performed three main analyses. First, we studied the association of EA-PGS with school performance in the full sample and compared the results to those from previous studies^6; 7^. In our sample, the maximum variance explained in school performance by EA-PGS was 8.4%. This was lower compared to the estimates reported in previous studies^6; 7^ (R^2^=14%). But subsequent analysis suggested that the relatively low R^2^ estimate in the full sample was partially due to the inclusion of those with psychiatric disorders (mainly ADHD and ASD) in the analysis. The maximum variance explained in school performance by EA-PGS raised to 10.6% when the analysis was performed only in controls. However, this is still slightly lower than the previous reports, which could be due to that we used SNP weights from the summary statistics from SSGAC that excluded 23andme samples^4^. But other factors such as environmental differences between our cohort and those involved in the previous studies are also likely to contribute as cohort to cohort genetic heterogeneity has been well documented in previous EA GWAS^4^. Nevertheless, our first analysis represents an independent replication of the strong association of EA-PGS with school performance.

Second, by using ADHD as reference group we were able to benchmark the effect sizes of EA-PGS deciles in controls. Our analysis showed that the school performance of controls in the lowest decile were comparable to the school performance of those with ADHD. Hence, it can be inferred that the impact of low EA-PGS on school performance might be comparable to the impact of ADHD (a condition that is strongly associated with learning difficulties) on school performance. However, we caution that the reference group in our analysis comprised of mainly high functioning ADHD individuals i.e. those who were able to go to school and sit for the exams. Should we have considered those who dropped out of school as well the impact of ADHD on school performance might have been stronger compared to the impact of low EA-PGS^35^ on school performance.

Nevertheless, the results we present here is to our knowledge the first demonstration that having a low EA-PGS is comparable to having learning difficulties. It is however important to be aware that these comparisons are meaningful only at group-levels and not at individual levels, which can be appreciated from the wide overlaps in the E1 distribution across the deciles.

Third, we demonstrate the EA-PGS’s association with school performance is weaker in individuals with ASD and ADHD than in the controls. Interestingly, the factor that reduces the strength of the association of EA-PGS with school performance in ADHD seems to be different from the one in ASD. The attenuation in the EA-PGS’s association with school performance in ADHD is mainly to due to a factor that is common between EA-PGS and SES. We speculate that this common factor is the genetic nurture i.e. the indirect genetic effects of the parents’ genotypes on offspring’s school performance mediated through the environment^10^. Recently it has been demonstrated that EA-PGS constructed using the nontransmitted alleles in the parents’ genome significantly correlate with offspring’s EA and this correlation is mediated through the rearing environment the parents create for their children^10^. The study reported that one third of the EA-PGS’s association with EA was mediated through genetic nurture. In line with this report, a recent study has demonstrated that EA-PGS’s association with EA is weaker in adopted individuals than in the nonadopted^36^. The variance explained in EA by EA-PGS in adopted was only half of that in nonadopted. This is because the adopted individuals’ do not share their genome with their foster parents and hence passive gene-environment correlation does not happen. In our study we found that both EA-PGS and parents’ education and employment statuses explained lower variance in school performance in those with ADHD than in the controls. This means that having educated and wealthy parents does not boost child’s school performance if the child has ADHD as much as it does if the child does not have ADHD. However, the underlying mechanism is not clear. We speculate that it could be due to a lack of the ADHD-child’s response to their parents’ nurturing effort, and as a result the passive gene-environment correlation (i.e. the correlation between parents’ nurturing behavior—which in turn correlates with their education levels—and EA associated genetic variants in the child shared with their parents) does not happen effectively. However, this is only speculation and needs confirmation through study designs ideal for probing passive gene-environment correlations, for example, by using genotyped parents-offspring trios, one can compare the association of parents’ EA-PGS constructed using untransmitted alleles with the offspring’s school performance between ADHD and control families.

Unlike ADHD, the variance difference between ASD and controls did not attenuate substantially after controlling for SES, which means that a lack of genetic nurture is unlikely to be the one that weakened the EA-PGS’s association with school performance in ASD. Other factors such as rare germline or de-novo copy number variations (CNVs) might have had a larger negative effect on school performance in ASD than controls in our sample, which in turn might have weakened the association of EA-PGS (which were based on common variants) with school performance^37^. In line with that, it has been shown that many of the CNVs particularly large deletions commonly seen in individuals with ASD have substantial negative effects on intelligence^38^. As we did not have CNV information for our sample, we were not able to test this hypothesis. We recommend future studies to evaluate both common and rare variants together when studying genetic effects on cognition in individuals with ASD.

Some of the limitations of our study are as follows. First, our study sample is biased toward those who were functional enough to go to school and sit for the exams. Hence, the results cannot be completely generalized, particularly the results from ASD. Second, not everyone in the psychiatric disorder groups received their diagnoses before the exams. However, it is not possible to know when the individuals received their first diagnosis as our register data contains only hospital-based records and do not contain information from local general practitioners and private practitioners. Hence, the first date of diagnosis cannot be considered as the age of disease onset and so we did not account for the date of diagnosis in our analysis. Third, the parents involved in our analysis are legal parents, but we did not know if they are biological parents as that information was not available. So, our interpretations in relation to genetic nurture are only speculative and need to be verified with genetic data from biological parent-offspring trios.

In summary, we have demonstrated that the genetic influences on school performance differ across individuals with psychiatric disorders and general population. Our results indicate that there might be a substantial heterogeneity in how genes and environment influence the school performance in individuals with psychiatric disorders and studying the same might offer novel insights into our understanding of the genetics of cognition.

## Subjects and Methods

### iPSYCH cohort

The study individuals are from iPSYCH^24^ and ANGI^25^, which are population-based Danish psychiatric case-cohorts recruited through Danish National registers^39; 40^. The iPSYCH cohort comprises of individuals diagnosed with at least one of five major psychiatric disorders (ADHD, ASD, MDD, SCZ and BPD) and a randomly selected controls without any of the five psychiatric disorders^24^. The ANGI cohort comprises of individuals diagnosed with AN recruited from United States, Sweden, Australia, and Denmark; only those recruited in Denmark through Danish registers were included in the current study^25^. ANGI participants were recruited and genotyped along with iPSYCH. Hence, all further descriptions about iPSYCH apply to ANGI as well. The iPSYCH cohort is a subsample of a large baseline cohort, which comprises of almost everyone born in Denmark between 1981 and 2005 who were alive on their first birthday and had a known mother (N=1,472,762)^24^. The phase 1 release, iPSYCH2012^24^, comprises of 77,369 individuals, ∼99% of whom were successfully genotyped (cases=51,101 and controls=27,605). Among them, around 60% of the individuals had information on school grades through Danish education registers. The rest of the individuals did not have information on school grades because they are either still young as of Dec 2016 or home schooled or dropped out of school. After sample quality control (QC) filtering, 30,982 individuals were included in the final analysis.

### School grades

The school grades were from exit exam (also called as ninth level exam or FP9) that marks the end of compulsory schooling in Denmark. The school grades of the iPSYCH individuals were obtained from Danish education register^41^. We chose three subjects namely Danish, English and mathematics, all of which have a compulsory exit exam hence available for most of the individuals. The types of grades available under each subject varied namely written, oral and grammar under Danish, oral under English and written and oral (for exams conducted until 2006) and problem solving (for exams conducted since 2007) under mathematics. The school grades data was processed following strict QC procedures that ensured minimal heterogeneity. Only those who were aged 14.5 to 17.5 years at the time of exam were included. Only the first attempt grades were included. Only those who sat for the exams in all three subjects in the same school year were included. After all the QC steps, a PCA was performed in the school grades and the first PC, named as E1, was extracted. As the mathematics grades differed between exams conducted between 2002-2006 and 2007-2016, we performed PCA separately in two datasets and then concatenated the PCs. The E1 explained substantial and equal proportion of variances across all the subjects hence indexing the overall school performance. All subsequent analysis we report here are based on E1. For more details on the school grades and the PCA of the school grades refer to Rajagopal et al 2020^22^.

### Psychiatric diagnoses

The cases in iPSYCH were identified based on the ICD-10 diagnoses recorded in the Danish Psychiatric Central Research Register^40^. The ICD-10 codes used for the six disorder were F90.0 for ADHD, F84.0, F84.1, F84.5, F84.8 and F84.9 for ASD, F32-F39 for MDD [Since 96% of the individuals had either F32 (depressive episode) or F33 (recurrent depressive disorder), we call it as MDD rather than as affective disorders], F30-31 for BPD, F20 for SCZ and F50.0 for AN. Individuals with ICD-10 diagnosis for mental retardation (F70-F79) were excluded.

### Ethical approvals

The iPSYCH project has been approved by the Danish Scientific Ethics Committee, the Danish Health Data Authority, the Danish Data Protection Agency and the Danish Neonatal Screening Biobank Steering Committee. For more details, refer to Pedersen et al^24^

### Genotyping and imputation

The iPSYCH individuals were genotyped using the DNA isolated from dried blood spots obtained from the Danish neonatal screening biobank (DNSB)^24^. The DNSB stores dried blood spots made using heel prick blood taken 4-7 days after birth for everyone born in Denmark since 1981^42^. The DNA was isolated from the blood spot and whole genome amplified^43^ in triplicates and genotyped using PsychChip v1.0 array. The genotyped markers were then phased using SHAPEIT^44^ followed by imputation using IMPUTE2^45^ with 1000 genomes phase-3 as the reference panel^46^. Standard QC procedures were followed both prior to as well as after imputation, which are described in detail elsewhere^30; 47^. After all QC steps, totally ∼6 million imputed variants with MAF>0.05 and INFO>0.80 were available for polygenic score construction.

### Relatedness and population stratification

Only unrelated individuals with European ancestries were studied. We excluded individuals of non-European ancestries because the EA-PGS was based on a GWAS^4^ performed only in Europeans and hence cannot be used to predict school grades in non-European sample. Relatedness was estimated using identity by descent (IBD) analysis using Plink v.1.90^48^. Pairs of individuals with PIHAT > 0.20 were considered as related and one of each pair was removed randomly. PCA of common variants was done in the unrelated individuals using the SNPRelate^49^ R package using a set of high-quality genetic variants. A subsample of the study individuals whose parents and paternal and maternal grandparents all born in Denmark were used as a reference group to identify population outliers. A five-dimensional ellipsoid based on the first five PCs of the reference group with a diameter of eight standard deviations was created and those who fell outside this ellipsoid were considered as non-Europeans and excluded from the analysis. The PCA was repeated after outlier removal and the first 10 PCs were included as covariates in the polygenic score association analysis.

### EA-PGS construction

EA-PGS was constructed using effect sizes from the recent large GWAS of educational attainment^4^. The publicly available summary statistics that excluded 23andme sample was used for this study (N= 766,345). The summary statistic was LD clumped using 1000 genomes^46^ EUR sample as reference to identify independent variants. Insertion and deletion variants, variants with ambiguous alleles (A/T and G/C) and low frequency variants (MAF < 0.05) were removed prior to clumping. The clumped summary statistics file is then used for polygenic scoring. In the target sample, only variants with INFO > 0.90 and missing rate < 1% were included. Polygenic scores were calculated using Plink v1.90^48^ at ten P value thresholds (5×10^−8^, 1×10^−6^, 0.0001, 0.001, 0.01, 0.05, 0.1, 0.2, 0.5, 1).

### EA-PGS and SES analysis

Both EA-PGS and SES association analyses were performed using linear regression adjusted for covariates. In the EA-PGS analysis, we included the following covariates: exam age, sex, first 10 ancestral PCs, genotyping waves, psychiatric diagnoses and school grades PCA group (since school grades PCA was performed separately in two datasets and the PCs are combined together, a dummy variable representing the two groups was included as a covariate). In the SES analysis, we included the following covariates: exam age, sex and school grades PCA group. In each case we performed two regressions: model 1, which included the main predictor (either EA-PGS or SES) and the covariates and model 2, which included only the covariates. The final R^2^, called as incremental R^2^, was obtained by subtracting the model 2 R^2^ from the model 1 R^2^. In order to statistically compare R^2^ between two regression analyses, we estimated standard error via bootstrapping. We repeated the regression analysis 1000 times with each time the samples chosen by bootstrapping with replacement and measured the incremental R^2^ each time. The SE is obtained by calculating SD of the resulting 1000 R^2^ values. The calculation of SE via bootstrapping was done using R package ‘boot’. Then we calculated Z score by dividing the R^2^ with SE and used the Z score for pairwise comparison using Z-test as described in Cheesman et al^36^.

### Socio-economic variables

We analyzed socio-economic variables namely mother’s education, father’s education, mother’s employment and father’s employment from the Danish registers. The parents’ education was extracted from Danish education registers and were analyzed as continuous variables with four levels namely primary education or no education (0), high school or vocational education (1), short cycle higher education or bachelor (3) and master or PhD (4). The parents’ employment information was as per the Danish labor market affiliations: retired (0), unemployed (1), employed-basic level (2), employed medium level (3) and employed top level (4). The descriptions of the five employment levels are provided in the Supplementary Table 11. The employment status was also analyzed as continuous variable. All these variables were as per the exam year i.e., the year when the student sat for the exam. If the variables were not available on the exam year, information from one or two years prior to exam year was extracted. We restricted our analyses to only those whose mother and father were alive on the exam year and those who resided with their mother and father in the same household on the exam year.

### Multiple testing corrections

The statistical significance in each of the analyses was evaluated after accounting for multiple testing using Bonferroni’s method. In all the PGS deciles-based analyses the P value threshold was set to 0.005 (0.05/10). In all the R^2^ comparisons between the psychiatric disorders and controls (both PGS and SES analyses), the P value threshold was set to 0.008 (0.05/6). In the R^2^ comparison across the SES quintiles, the P value threshold was set to 0.01 (0.05/5).

### Supplemental Data

Figure 2 has one figure supplement; figure 5 has one figure supplement; summary statistics from all the statistical analyses were provided as supplementary tables (1-11) in an excel file.

## Acknowledgements

1. The iPSYCH project is funded by the Lundbeck Foundation (grant numbers R102-A9118, R155-2014-1724 and R248-2017-2003) and the universities and university hospitals of Aarhus and Copenhagen. The Danish National Biobank resource was supported by the Novo Nordisk Foundation. Data handling and analysis was supported by NIMH (1U01MH109514-01 to ADB). High-performance computer capacity for handling and statistical analysis of iPSYCH data on the GenomeDK HPC facility was provided by the Center for Genomics and Personalized Medicine, Aarhus University and Central Region Denmark, and Centre for Integrative Sequencing, iSEQ, Aarhus University (grant to ADB).
2. The Anorexia Nervosa Genetics Initiative (ANGI) was an initiative of the Klarman Family Foundation.
3. The PhD fellowship of V.M.R was fully funded by the Graduate School of Health, Aarhus University, Aarhus, Denmark.
4. We thank Rosa Cheesman (King’s College London) and Jonathan R. I. Coleman (King’s College London) for providing scripts to estimate standard errors for R2 using R boot package and to perform Z test.
5. We thank Varun Warrier (University of Cambridge) for the useful discussions and thoughtful comments on the manuscript.

## Data and code availability

The EA3 summary statistics used for polygenic scoring is publicly available at https://www.thessgac.org/. The summary statistics related to the analyses reported in this paper are available in the supplementary tables.

The code supporting the current study have not been deposited in a public repository because they were written within secured servers: GenomeDK (genome.au.dk) and Denmark statistics (DST; dst.dk) and were tailored for the iPSYCH data. However, code related to any specific analysis can be exported from servers through proper channels if necessary.

---

**Figure 2—figure supplement 1.**
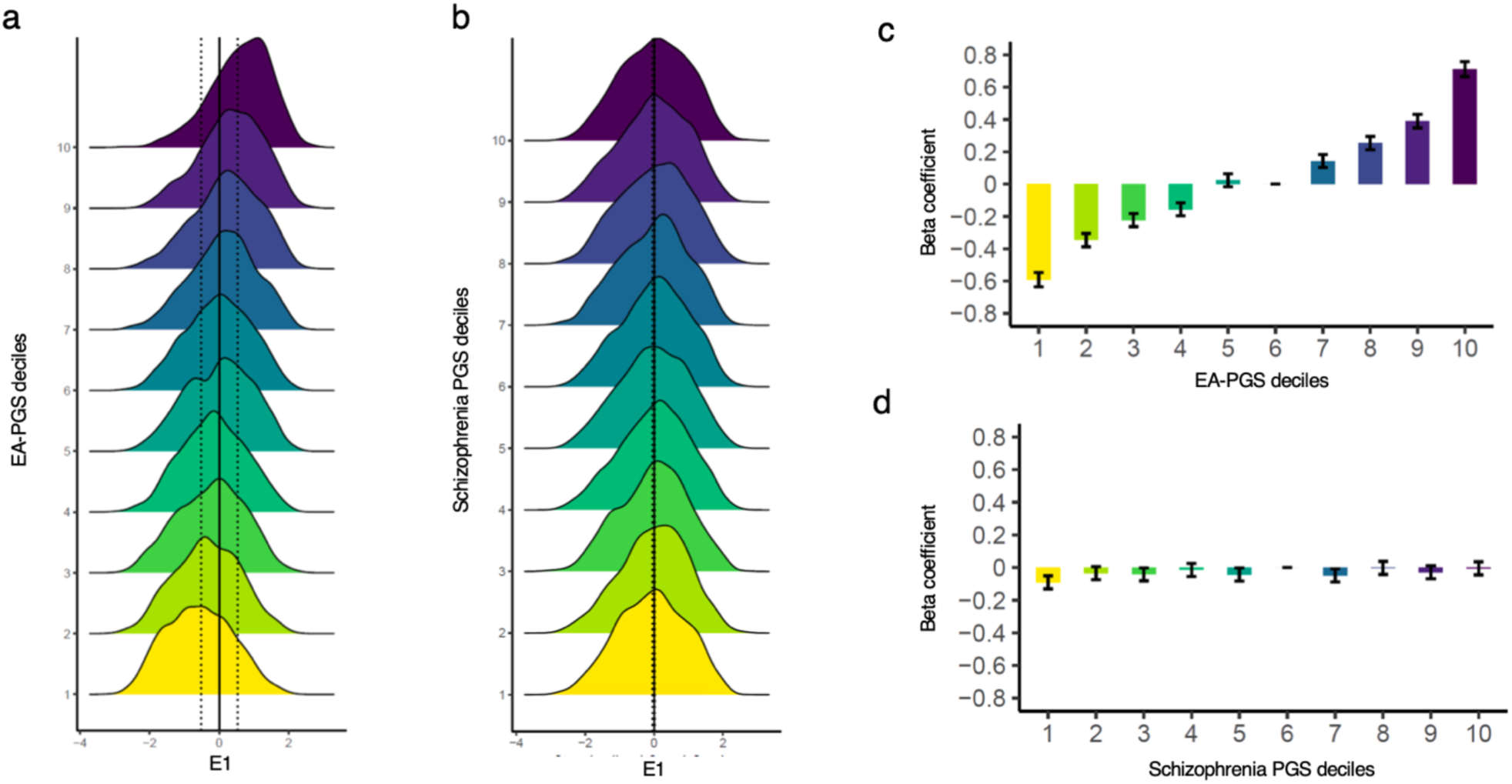
Distribution of E1 across PGS deciles (in controls). The description here is same as the main figure 2, except that here only the control individuals (N=12,487) were analysed.

**Figure 5—figure supplement 1.**
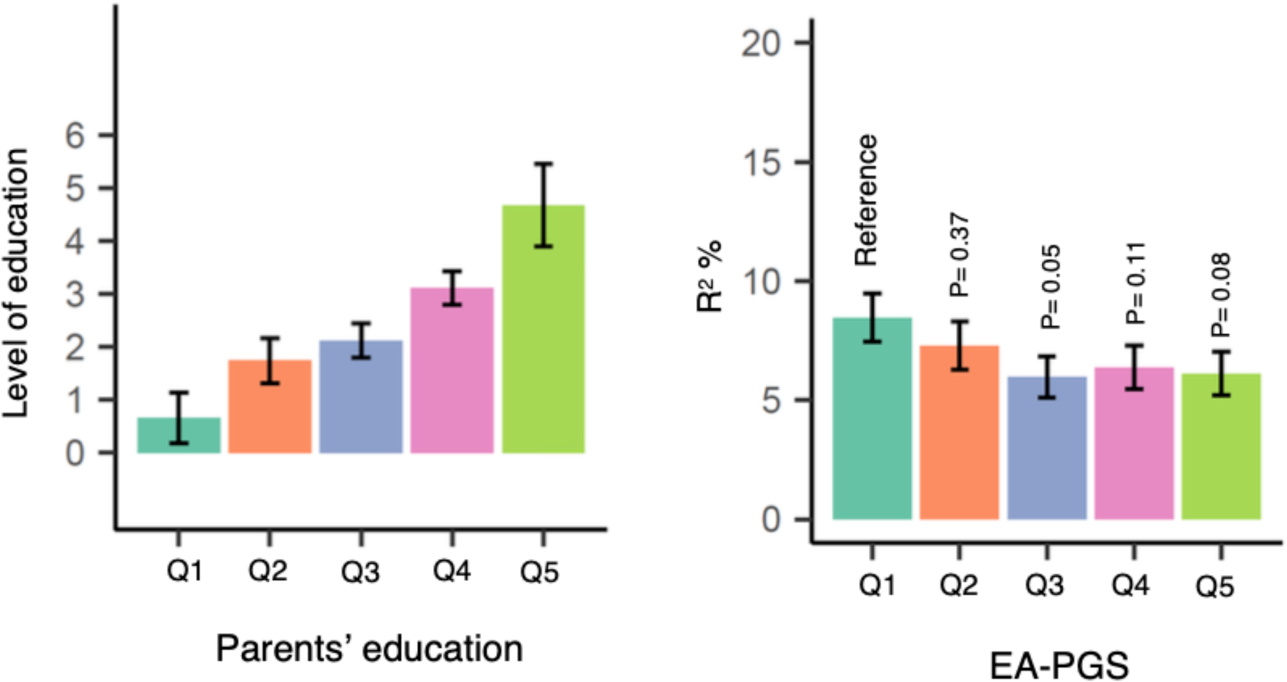
Association of EA-PGS with E1 across SES quintiles. The controls were divided in to quintiles based on their parents’ education (father’s and mother’s education combined) and the association of EA-PGS with E1 was tested in each of the quintiles. **a**, Mean SES in each of the quintiles; error bars represent standard errors **b**, Variance explained in E1 by EA-PGS in each of the quintiles. The R^2^ in quintiles 2-5 were tested if they are statistically different from the R^2^ in quintile 1 using Z test and the P values are provided in the plot.

